# Genomic landscape of reproductive isolation in *Lucania* killifish: The role of sex chromosomes and salinity

**DOI:** 10.1101/831867

**Authors:** Emma L. Berdan, Rebecca C. Fuller, Genevieve M. Kozak

## Abstract

Understanding how speciation occurs and how reproductive barriers contribute to population structure at a genomic scale requires elucidating the genetic architecture of reproductive isolating barriers. In particular, it is crucial to determine if loci underlying reproductive isolation are genetically linked or if they are located on sex chromosomes, which have unique inheritance and population genetic properties. Bluefin killifish (*Lucania goodei*) and rainwater killifish (*L. parva*) are closely related species that have diverged across a salinity gradient and are reproductively isolated by assortative mating, hybrid male infertility, viability of hybrid offspring at high salinities, as well as reduced overall fitness of F2 offspring and backcrosses to *L. goodei*. We conducted QTL mapping in backcrosses between *L. parva* and *L. goodei* to determine the genetic architecture of sex determination, mate attractiveness, fertility, and salinity tolerance. We find that the sex locus appears to be male determining and located on a chromosome that has undergone a Robertsonian fusion in *L. parva* relative to *L. goodei*. We find that the sex locus on the fused chromosome is involved in several genomic incompatibilities, which affect the survival of backcrossed offspring. Among the backcrossed offspring that survived to adulthood, we find that one QTL for male attractiveness to *L. goodei* females is closely linked to this sex locus on chromosome 1. Males homozygous for *L. goodei* alleles at the sex locus laid more eggs with *L. goodei* females. QTL associated with salinity tolerance were spread across the genome but did not tend to co-localize with reproductive isolation. Thus, speciation in this system appears to be driven by reinforcement and indirect selection against hybrids rather than direct natural selection for salinity tolerance. Our work adds to growing evidence that sex chromosome evolution may contribute to speciation.

## INTRODUCTION

Progress towards speciation can depend on extrinsic interactions of populations with their environment and intrinsic genomic architecture that separately or together cause a reduction in gene flow (Campbell *et al.*, 2018). Gene flow and recombination directly oppose divergence and speciation because they homogenize allelic combinations that are unique to each population (Felsenstein, 1981; Butlin, 2005). Adaptation to abiotic and biotic features of the environment can lead to phenotypic changes among populations, causing reductions in mating rate or hybrid viability that reduce the probability of gene flow (Coyne & Orr, 2004; Schluter, 2009; Nosil, 2012). Rearrangements in chromosomal structure reduce recombination by suppressing it between homologous chromosomes with different arrangements. If the genes that underlie reproductive isolation and/or ecological divergence are present in regions of low recombination, then they are protected from gene flow even when hybridization occurs, making genome divergence and ultimately speciation much more likely (Noor *et al*., 2001; Kirkpatrick & Barton, 2006; Hoffmann & Rieseberg 2008, Faria & Navarro, 2010; Lowry & Willis, 2010; Wellenreuther & Bernatchez, 2018; Charlesworth & Barton, 2018; Wellenreuther *et al*., 2019). Sex-determining loci can also function to reduce recombination across a chromosome when heterozygous, which can lead to faster genomic divergence on sex chromosomes (Meisel & Connallon, 2013; Sackton *et al.*, 2014). The relative roles of external forces and internal architectural features in divergence is an area of active speciation genomics research (Campbell *et al.*, 2018).

Genomic studies that map traits relevant to environmental features and reproductive isolation are key to understanding the relative roles of extrinsic and intrinsic forces in speciation. As the process of speciation involves multiple reproductive isolating barriers that reduce gene flow among incipient species, it is important to study how these barriers build up, become associated with one another in the genome, and potentially generate emergent reproductive isolation when coincident (Butlin & Smadja, 2018). If a chromosomal rearrangement has facilitated ecological divergence, the expectation would be that ecologically important traits map to the rearranged region. Linkage of multiple ecological traits can drive the spread of a rearrangement in theoretical models (Kirkpatrick & Barton, 2006). Chromosomal rearrangements may also be expected to link multiple forms of reproductive isolation, such as loci causing assortative mating with those contributing to hybrid incompatibilities (Trickett & Butlin, 1994; Dagilis & Kirkpatrick, 2016). Due to reduced recombination and increased genomic divergence, sex chromosomes also contribute substantially to speciation, harboring more reproductive isolating loci than other chromosomes (Coyne, 1992; Turelli & Orr 2000; Presgraves, 2008; 2018). This is often referred to as the “large X effect” although it occurs on all types of sex chromosomes (Z: Dopman et al. 2004; W: Saether *et al*., 2007; neo-Y: Kitano *et al.,* 2009).

One of the key extrinsic features that contributes to speciation in marine environments is adaptation to salinity (Lee & Bell, 1999; Hrbek & Meyer, 2003; Huyse *et al.*, 2004; Whitehead, 2010; Betancur *et al.,* 2015). Environmental salinity requires complex physiological adaptation because in high salinity environments, organisms are subject to ion influxes and loss of water from tissues. Conversely in low salinity environments, fluxes of water into tissues and loss of ions to the environment occurs (Evans *et al.,* 2005; Evans, 2008). This complex adaptation causes divergence in many tissues and can lead to speciation as a direct consequence of adaptation to salinity (Taylor, 1999; Seehausen & Wagner, 2014). Previous work suggests the genomic basis of this important physiological trait may be dispersed across the genome. For example, in Atlantic cod, adaptation to salinity was associated with outlier loci on 11 out of 23 linkage groups (48%) (Berg *et al.,* 2015). However, it remains unknown if salinity tolerance loci might be genetically linked to traits directly related to reproductive isolation, particularly in species that have diverged along a salinity gradient.

Here we map salinity tolerance and reproductive isolation across the genome relative to internal features, including a chromosomal fusion and the sex locus, in two hybridizing species of *Lucania* killifish. *Lucania goodei* and *L. parva* are recently diverged sister species (Duggins *et al.,* 1983; Whitehead, 2010) that differ radically in their salinity tolerance*. Lucania goodei* is found primarily in freshwater sites (restricted mainly to Florida and southern Georgia), while *Lucania parva* can be found in fresh, brackish, and marine habitats as far west as central Mexico and as far north as Massachusetts (Lee, 1980). Differential adaptation to salinity between the two species is present at multiple life stages (Dunson & Travis, 1991; Fuller *et al*., 2007, Fuller, 2008). Hybrids between *L. parva* and *L. goodei* can be found in the wild (Hubbs *et al.,* 1943), but multiple reproductive isolating barriers exist. Hybrid sons of *L. parva* females and *L. goodei* males have reduced fertility, there is reduced viability of hybrid offspring at high salinities, and reduced overall fitness of F2 offspring and backcrosses to *L. goodei* (Fuller *et al.*, 2007; Fuller 2008). Assortative mating due to male and female preferences also exists between the two species (Fuller *et al*., 2007; Berdan & Fuller 2012; Kozak *et al.* 2015; St. John & Fuller, 2019). Several salinity and fertility related genes show divergence among *L. parva and L. goodei* (Kozak *et al.*, 2014). In *L. parva*, a Robertsonian chromosomal fusion has occurred and two acentric chromosomes have been fused into a single metacentric one (Berdan *et al.*, 2014). The sex determining locus is currently unmapped in these species.

We genetically mapped the sex determining locus, salinity tolerance, behavioral isolation (female preference and male attractiveness/preference for each species), and intrinsic postzygotic isolation (reduced hybrid survival and reduced male fertility) using crosses between these species. We wanted to determine if these traits mapped to the same area of the genome and, in particular, if the traits are linked to the chromosomal fusion or the sex locus. To do this, we created a series of backcrossed hybrids (backcrossed to *L. goodei*), phenotyped the backcrossed offspring for salinity tolerance, female mating preferences, male attractiveness/preference, and male fitness, and genotyped the offspring at 4,545 SNPs for map construction and QTL mapping.

## METHODS

### QTL Mapping Cross

For the QTL mapping of reproductive isolating traits, we created backcrosses to *L. goodei.* The parental adult *L. goodei* and *L. parva* were collected from a sympatric population at the Oklawaha River at the Boat Ramp at Delk’s Bluff near Ocala (Marion County, Florida). We subsequently had difficulty obtaining enough *L. parva* from this site to use as stimulus animals in our behavioral assays (see below), so we also obtained stimulus animals from another sympatric population on the Wakulla River (Wakulla County, Florida).

All individuals were collected using dip nets and seines between 2009-2011. Animals were transported back to the University of Illinois where they were housed by population in 76-liter (20 gallon) aquaria, 110-liter (29 gallon) aquaria, and 568-liter stock tanks. In all experiments, our freshwater source was dechlorinated city water treated with Start Right (Jungle Laboratories, Cibolo, TX). Fish were fed ad lib daily with frozen brine shrimp. Lights were maintained on a 14L:10D cycle.

### Backcrosses to L. goodei

We created a series of backcrossed hybrid offspring (backcrossed to *L. goodei*) that we used for the experiments. In September 2009, we set up our F1 crosses. We performed F1 crosses in both directions (F1 – *L. goodei* ♀ X *L. parva* ♂, F1r – *L. parva* ♀ X *L. goodei* ♂) using fish that occurred in sympatry at the Boat Ramp at Delk’s Bluff. We originally set up 5 replicates of each cross. Each pair of fish was placed in a 38-liter aquarium (10 gallon) with four yarn mops that served as spawning substrate. Tanks were checked for eggs every 2-3 days. In November 2009, we added 7 additional replicates: 3 F1 crosses and 4 F1r crosses. Egg checking continued through April 2010. Eggs were placed in tubs of freshwater and treated with dilute methylene blue (an anti-fungal agent). After hatching, fry were fed with newly hatched *Artemia salina*. We recorded the number of eggs that hatched and the number of fry that survived to one month. At one month of age, fry were put into 110-liter (29 gallon) aquaria where they were raised to adulthood. We used the adult F1 offspring to create backcrosses to *L. goodei* in July - August 2010. All of the *L. goodei* used in the creation of the backcrosses were from the Delk’s Bluff population. We created all four types of backcrosses: BC1-F1 ♀ X *L. goodei* ♂, BC2- *L. goodei* ♀ X F1 ♂, BC3-F1r ♀ X *L. goodei* ♂, and BC4- *L. goodei* ♀ X F1r ♂. Each pair of fish was placed in a 38-liter aquarium (10 gallon) with four yarn mops that served as spawning substrate. Tanks were checked for eggs every 2-3 days. A portion of the eggs were used in salinity tolerance assays and the remainder were raised to adulthood for use in mate choice assays. Husbandry was identical to that described above for the F1 offspring.

### Salinity tolerance

For the salinity tolerance assay, we divided clutches of eggs from backcrosses between fresh water and salt water. Half of the eggs were placed in fresh water (0.2 ppt), and the other half were placed in salt water (15 ppt). For the freshwater treatment, eggs were placed in 177 mL (6 ounce) tubs of fresh water (dechlorinated city water) treated with methylene blue (anti-fungal agent). For the saltwater treatment, eggs were placed in tubs containing water at 15 ppt and treated with methylene blue. Our saltwater source was reverse osmosis water from a 4-stage barracuda RO/DI unit (Aqua Engineering and Equipment, Winter Park, Florida) to which we added Instant Ocean® Sea Salt (Spectrum Brands, Atlanta, GA) to achieve the desired salinity. Salinity was verified with an YSI-63 salinity meter (YSI Inc., Yellow Springs, OH). After hatching, fry were fed with newly hatched *Artemia salina*. All fry were raised to one month of age and euthanized with MS-222 (Argent Chemical Laboratories, Redgemont, WA). Offspring were stored in ethanol at −20° C until subsequent DNA extraction. We recorded the number of eggs that hatched and the number of fry that survived to one month.

### Behavioral isolation

We assayed adult backcrossed female mating preferences in June and July of 2011. We used a no-choice mating assay which has been used successfully in previous studies of behavioral isolation in *Lucania* (Fuller *et al.*, 2007; Berdan & Fuller 2012; Kozak et al. 2012; St. John & Fuller, 2019). Backcrossed females were placed in a 38-liter (10 gallon) aquarium with a stimulus male; either a male *L. goodei* or a male *L. parva*. All of the stimulus males came from the Delk’s Bluff populations. All tanks were provided with four yarn mops that served as spawning substrate. This resulted in 8 experimental treatments (four types of females and two types of males). We endeavored to have 5 replicates of each but actual replication varied depending on the availability of fish. We conducted the following number of replicates: assays with *L. goodei* males BC1 = 4, BC2 = 2, BC3 = 6, BC 4 = 3; assays with *L. parva* males BC1 = 4, BC2 = 1, BC3 = 4, BC4 = 3; resulting in 27 females total. All females were only tested with one male. These tanks were checked for eggs every 2^nd^ day for 21 days. From these data, probability of mating, latency to mate and average egg production was calculated. At the end of the experiment, all females were euthanized with MS-222 and stored in ethanol at −20° C.

We assayed male backcrossed offspring for male preference/attractiveness in August and September of 2011. Here, we also used a no-choice mating assay. Backcrossed male offspring were placed in a 38-liter (10 gallon) aquarium with a stimulus female: either a female *L. goodei* or a female *L. parva*. We originally planned for all of the stimulus females to come from the Delk’s Bluff population. However, low abundance of *L. parva* at that site in August 2011 rendered this impossible. We created as many tanks as possible using Delk’s females (12 tanks: 6 with *L. goodei* females, and 6 with *L. parva* females), and we used female *L. goodei* and *L. parva* from the Wakulla River population for the remaining 28 tanks. Delk’s Bluff and Wakulla River are both sympatric freshwater sites. We endeavored to create equal replication for each female species by male backcross combination, but actual replication varied depending on availability of fish. We conducted the following number of replicates: assays with Delk’s Bluff *L. goodei* females BC1 = 3, BC2 = 0, BC3 = 0, BC4 = 2; assays with Wakulla River *L. goodei* females BC1 = 7, BC2 = 1, BC3 = 12, BC4 = 4; assays with Delk’s Bluff *L. parva* females BC1 = 4, BC2 = 0, BC3 = 1, BC4 = 2; assays with Wakulla River *L. parva* females BC1 = 8, BC2 = 1, BC3 = 13, BC4 = 4. Overall 29 males were tested with both *L. goodei* and *L. parva* females and 4 were tested only with *L. parva* females. Males tested with both females (random order) were paired with a given female for 20 days, and then subsequently paired with a stimulus female of the opposite species (but from the same population). This resulted in 33 males tested in total (33 with *L. parva*; 29 with *L. goodei*). Tanks were checked every other day for eggs. Probability of mating (yes or no), latency to mate and egg production data were calculated and served as indices of male attractiveness/female choice. After mating trails, males were subsequently euthanized with MS-222 and stored in ethanol at −20° C.

### Reduced male reproductive success

Previous work on *Lucania* indicates that a large genetic incompatibility is segregating between the two species that results in some hybrid males having drastically reduced fitness (Fuller 2008). Nearly half of the offspring from male hybrid F1r (*L. parva* female × *L. goodei* male) die during the first few days of development compared to those from male F1 hybrids (*L. goodei* female × *L. parva* male). We assayed both the fertilization success and the survival of eggs spawned by the various backcross males. We checked all collected eggs under the microscope to assess fertilization. We considered eggs that were already dead upon collection to be unfertilized. We saved the fertilized eggs and measured their survival until hatching. We surveyed a total of 23 males for which we have two measures of male reproductive success: fertilization success and survival to hatching.

### SNP genotyping and linkage map construction

DNA was extracted using a modified version of the PureGene (Gentra Systems, www.gentra.com) extraction protocol over four days. On the first day, tissue samples were placed in 600 μl of cell lysis solution (0.1 M Tris, 0.077 M EDTA, and 0.0035 M SDS) with 3 μl of Proteinase K (20 mg/ml). The samples were vortexed and kept at 65° C overnight. On the second day, 200 μl of protein precipitation solution (Qiagen, Valencia, CA) was added to each, and the samples were vortexed and then stored at 4° C overnight. On day three, the samples were centrifuged at 12.6 rpm for 5 minutes. For each sample, the supernatant was removed leaving behind the protein pellet. Six hundred μl of isopropanol was added and the sample was kept at −20°C overnight. On the final day, the sample was centrifuged at 12.6 rpm for 4 minutes to precipitate the DNA. The supernatant was removed and 600 μl of 70% ethanol was added. The sample was vortexed and then centrifuged again. The ethanol was removed and the pellet was allowed to dry and then rehydrated with 30 μl of TE. Sample concentration and quality were verified using a Nanodrop spectrophotometer. DNA was extracted from 173 offspring from the salinity tolerance assay (61 freshwater, 84 saltwater), 33 males from the male behavioral isolation and intrinsic isolation assays, and 27 females from the female behavioral isolation assay. Samples were diluted to a concentration of 75 ng/μl prior to genotyping.

Species-specific SNPs were designed for the Illumina Infinium assay as described in Berdan *et al.* (2014). DNA samples were genotyped at all SNPs using an Illumina Infinium Bead Chip custom designed for *Lucania*. Bead chips were scanned using the iScan System (Illumina) at the Keck Center for Comparative and Functional Genomics at the University of Illinois. Raw data from the Infinium assay were changed to genotype calls using Illumina GenomeStudio software v2011.1. Cluster positioning was done automatically for species-specific SNPs. Afterwards cluster positioning was checked manually and minor adjustments were made to optimize genotype calls. The no-call threshold was set to 0.15 and genotype calls were exported as spreadsheets.

A hybrid linkage map was constructed from F1 hybrid parents using species-specific SNPs in Joinmap 4.0 (Li *et al.,* 2008) following methods used for constructing *L.parva* and *L. goodei* maps as described in Berdan *et al.* (2014).

### QTL mapping

All QTL mapping and other loci association tests were done in R v.3.5 (R Core Team, 2018). The distributions of all mapped phenotypes are shown in Supplemental Figure 1 and 2. We performed QTL analyses separately for all traits. Traits involved in behavioral isolation were separated by species as loci underlying *L. parva* species recognition might be different than traits underlying *L. goodei* species recognition. For each species, we analyzed two measures of behavioral isolation separately: probability of mating and egg production. In the crosses, individuals tended to mated quickly or not at all (see Figure S2), therefore we mapped probability of mating (whether or not mating occurred over 20 days) as opposed to latency to mate. Egg production was measured as the average number of eggs produced per day. These were measured for male backcrossed individuals and female backcrossed individuals separately. Thus, we had 8 traits that we mapped for behavioral isolation: male preference/attractiveness to each species as evidenced by egg production and latency to mate (4 traits), female preference/attractiveness to each species as evidenced by egg production and latency to mate (4 traits). For each of these traits, the QTL mapping was done in rQTL using the hybrid linkage map and scanone with standard mapping (Broman & Sen, 2009). Probability of mating used a binary model. We calculated the significance of LOD scores using 500 permutations and the 95% Bayesian credible interval for any significant QTL identified. We also looked for multiple interacting QTL using the scantwo function, but did not detect any significant QTL. This scantwo analysis may have been limited in power due to sample size.

### Gametic disequilibrium analyses – Interactions Among Loci

The goal here was to determine whether backcrossed offspring differed in their probability of survival due to interactions among genotypes located on different linkage groups. Incompatible loci should generate distortions in genotype frequencies in surviving backcrossed individuals. To do this, we tested for non-random patterns of genotypes, using a chi-squared analysis. We only included backcrossed offspring that had been raised in fresh water (61 individuals) to avoid the distorting effects of differential survival in salt water. We considered offspring who were raised until one month of age (excluding adult backcrossed offspring had little effect on the results). Along a given linkage group, many of the markers were in complete linkage, so we used one representative marker from each set in complete linkage. We also only considered patterns among loci located on different linkage groups. Hence, we did not test for interactions among loci on the same linkage group. We performed a total of 10,675 tests. For each test, we measured Chi-squared, the associated p-value, and the frequencies of the four combinations of genotype (homozygous at both locus 1 and 2, heterozygous at both locus 1 and 2, homozygous at locus 1/heterozygous at locus 2, and vice versa). We corrected for multiple testing by using the Benjamini and Hochberg false discovery rate (1995) method as implemented in R with ‘p.adjust’ statement.

### Salinity tolerance genotype testing

We sought to determine the location of QTL associated with salinity tolerance. To do this, we compared the frequency of the different genotypes across the genome among offspring raised in freshwater and saltwater. Survival was lower among offspring raised in salt water (20.9%) than in fresh water (39.4%). Previous work indicates that juveniles of both *L. goodei* and *L. parva* survive well in hard, fresh water. We therefore used the frequency of the SNP genotypes among the 61 freshwater offspring as the expected frequency and asked whether the frequencies in saltwater differed using the binomial test. We corrected for multiple testing by using the Benjamini and Hochberg (1995) method as implemented in R with ‘p.adjust’ statement.

### Mapping of the sex determining locus

Karyotypes of both *L. goodei* and *L. parva* suggested that the sex chromosomes were homogametic (Uyeno & Miller 1971; Berdan *et al.*, 2014). Therefore, we evaluated the possibility of a male determining locus as well as a female determining locus. To search for markers linked to the sex determining locus, we generated predictions about species-specific markers when different types of F1 hybrids were backcrossed to *L. goodei* (Table S1). For instance, if the sex locus is male determining (Y-like), then hybrid male offspring of an *L. parva* female and an *L. goodei* male (*L. parva* ♀ X *L. goodei* ♂) should pass on an *L. goodei* allele to male offspring and an *L. parva* allele to female offspring. When backcrossed to *L. goodei*, we expect female offspring to be heterozygous and male offspring to be homozygous for *L. goodei* alleles for loci linked to the sex locus. If the sex locus is female determining (W-like), then we expect hybrid females to pass on an *L. parva* allele only to female offspring (Table S1). We used the QTL mapping cross (backcrosses into *L. goodei*) to test predictions concerning the nature of sex determination (X-Y versus Z-W) and map the location of the sex determining locus. In addition, we used animals from two other crosses (one cross between *L. goodei* and *L. parva* and another between *L. parva* populations) from another study to independently map the location of the sex determining locus. In all crosses, we tested for an association between alleles and our predictions using rQTL with the predicted sex-linked loci coded as a binary phenotype (0 for homozygous, 1 for heterozygous). We used scanone with a binary model to calculate LOD scores, the significance using 500 permutations and the 95% Bayesian credible interval.

To map the male determining loci more finely, we used backcrossed offspring from another study. In this study, we created another set of hybrid offspring between the two species. Here, we used two allopatric populations: *L. goodei* from Blue Springs in the Suwanee/Santa Fe River (Florida) and *L. parva* from Indian River Lagoon (Atlantic Ocean, Florida). Collection methods and animal husbandry were identical to those described above for the QTL crosses. We used these offspring from backcrosses between these populations to independently verify the location of the sex-determination locus. In this study, we generated all possible backcrosses to both *L. goodei* and *L. parva* using both F1 and F1r hybrids parents. We genotyped 50 backcross offspring (32 from backcrosses to *L. goodei*, 18 from backcrosses to *L. parva*). For this analysis, we only considered species-specific SNPs (1030 SNPs; 353 of which had a position on the maps). We separately used the *L. goodei* and *L. parva* maps (Berdan *et al.* 2014) for mapping to see if this influenced the position of the sex locus. Table S2 shows the predicted genotypes for males and females for backcrosses to both *L. goodei* and *L. parva.*

We also created a series of hybrid crosses between two *L. parva* populations (Indian River, Florida and Pecos River, Texas). We created hybrids in both directions and created all backcross types. Collection methods and animal husbandry were identical to those described above for the QTL crosses. We genotyped 35 hybrid backcrossed individuals. We genotyped 14 offspring (7 females, 7 males) from F1 males (Indian River ♀ × Pecos ♂) and 21 offspring (11 females, 10 males) from F1r males (Pecos ♀ × Indian River ♂). We filtered SNP data and only used alleles that were fixed between Indian River and Pecos (Kozak *et al*., 2014) for a total of 1048 SNPs (821 of which had a position on the *L. parva* map). We mapped the sex-locus using the *L. parva* linkage map. Again, we tested the genotypes for the expected ratios of heterozygotes/homozygotes in males and females from backcrosses to each population (Table S3).

All plots were made in R using rQTL, ggplot2 (Wickham, 2017) and LinkageMapView packages (github.com/louellette/LinkageMapView).

## RESULTS

### Sex-determining locus

In both the *L. parva* map and the hybrid map, linkage group 1 represents a fusion of two linkage groups (1A and 1B) from *L. goodei*. All maps had 22 additional linkage groups and are numbered based on synteny (see Berdan *et al.,* 2014). Using the hybrid linkage map from the QTL cross, no female sex determining locus was found with all LOD < 1.32 (p > 0.53; N = 44 informative individuals). In contrast, we found evidence for a single male-determining sex locus on chromosome 1 at 0 cM near marker 05836 (LOD = 3.35, p=0.014, 95% Bayesian Credible Interval 0-12 cM). Using Indian River *L. parva* and Blue Springs *L. goodei* hybrids backcrossed to *L. goodei* and *L. parva* with the *L. goodei* linkage map, the male sex determining locus was located on chromosome 1A at 2 cM between markers 13121 and 14413 (LOD = 5.21, 95% Bayesian Credible Interval 0.5-3 cM; Figure 1A). Using these same data and the *L. parva* map, the sex locus was on chromosome 1 at 10.5 cM near marker 13005 (LOD = 6.82, p < 0.001, 95% Bayesian Credible Interval 9-11 cM; Figure 1B). Using crosses among *L. parva* populations (Indian River and Pecos River) backcrossed males and the *L. parva* map, the QTL for the sex determining loci was located on chromosome 1 at marker 11321 at 20.81 cM (LOD = 7.41, p < 0.001, 95% Bayesian Credible Interval 13-44 cM; N = 36). Thus, the sex determining locus consistently maps to the chromosome 1A portion of the fused chromosome. Among the *L. parva* within species/between population crosses, much of the chromosome appears to be in tight linkage disequilibrium with the sex loci (Figure 1C).

**Figure 1.**
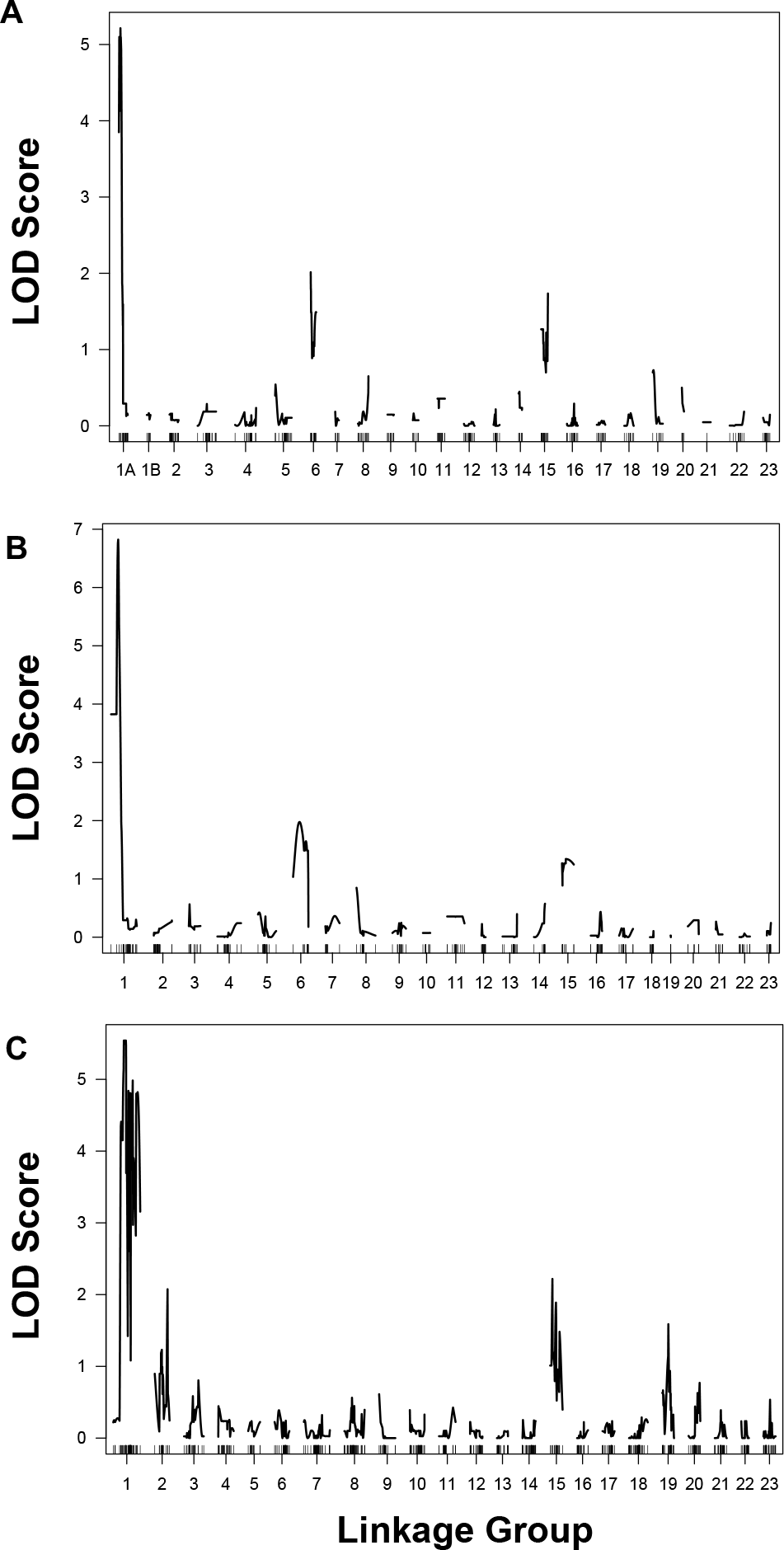
Sex determining locus linkage mapping. *L. parva* and *L. goodei* backcrosses using (A) *L. goodei* map and (B) *L. parva* map. (C) Sex determining locus in *L. parva* between population crosses using *L. parva* map. Although individual LOD score thresholds vary, LOD score > 3 is equivalent to p < 0.05.

### Gametic Disequilibrium – Interactions Among Loci

The chromosomal fusion was implicated in genetic incompatibilities. The backcrossed offspring who survived to one month of age were a non-random subset that had favorable combinations of alleles at different loci. Twenty-six of 10,675 tests for interactions among genotypes at loci on different linkage groups remained significant even after correcting for multiple tests. Table 2 lists these markers and the linkage groups on which they are found. While there were 26 significant interactions, these involved loci on only five pairs of linkage groups. There were multiple significant interactions involving loci on linkage group 1 and both linkage groups 13 and 16. One interaction between linkage group 1 and linkage group 13 involved a marker very close to the sex determination region (marker 13005). There were also significant interactions between linkage groups 13 and 16, linkage groups 21 and 22, and linkage groups 23 and 2. The interaction between linkage group 21 and 22 is interesting because it involves markers that mapped to linkage group 21 in one species and linkage group 22 in the other (a putative translocation: Berdan *et al.* 2014). All of these interactions among loci involved an over-representation of offspring that were either homozygous for the *L. goodei* specific marker at both loci or were heterozygous at both loci. Individuals that were homozygous at one locus, but heterozygous at another were either absent or under-represented. Supplemental table 4 contains all of the tests.

**Table 2.**
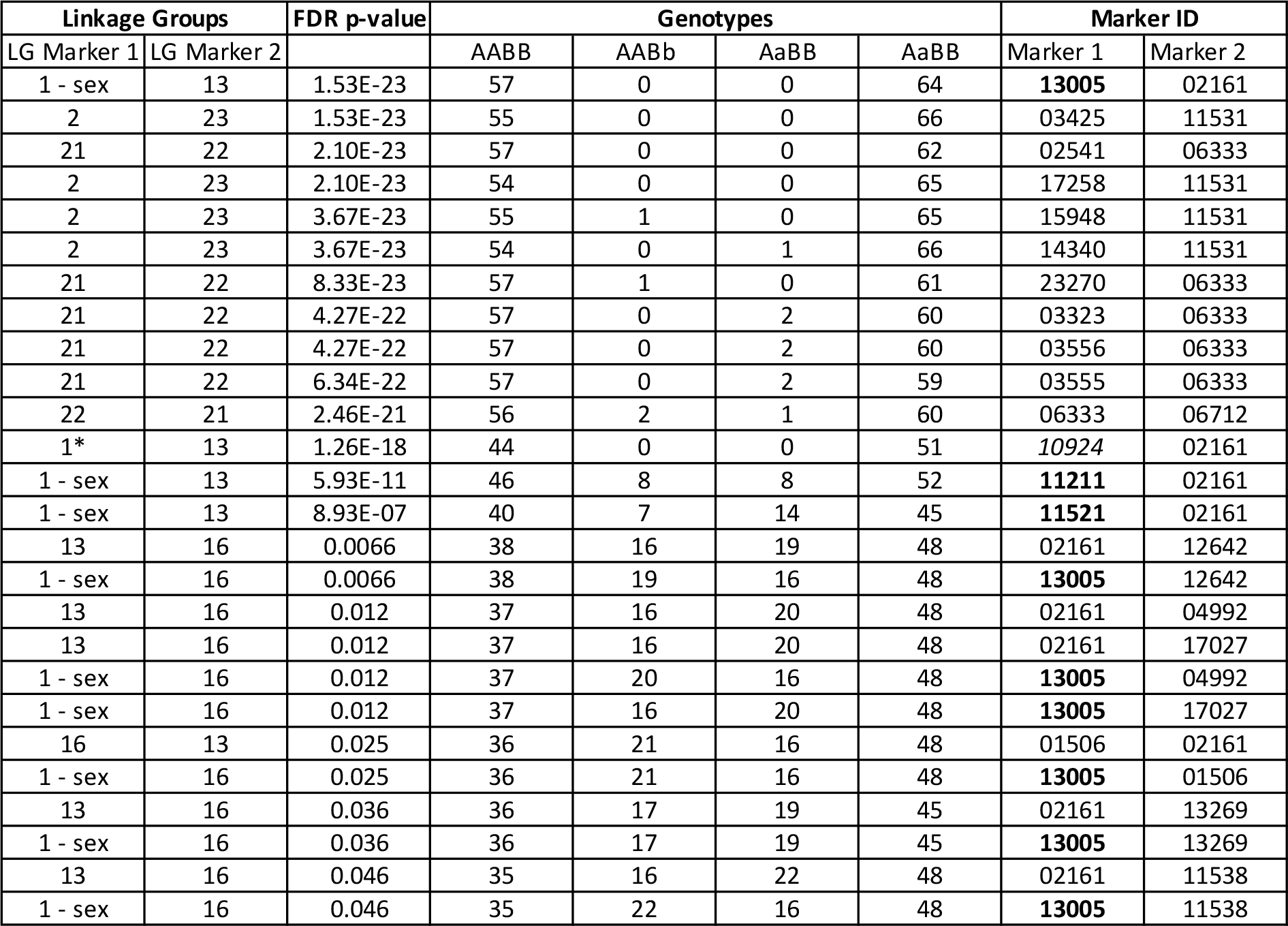
Genotypes between markers on different linkage groups with significant frequency distortion. Genotypes refers to the number of individuals that are homozygous for the *L. goodei* marker at both loci (AABB), are heterozygous at both loci (AaBb), or are homozygous at one locus but heterozygous at another (AABb and AaBB). 1-sex indicates a marker on linkage group1 located within the sex determining region (marker ID in bold); 1* indicates a marker on linkage group 1 that is adjacent to the sex determining region (marker ID in italics).

### Salinity tolerance

Survival in salt water was approximately half of that in in fresh water (salt water = 20.9%; fresh water = 39.4%). Backcross survival to one month of age in saltwater was affected by genotype. We compared the proportion of homozygous (*L. goodei*) and hybrid genotypes at each marker between fresh and saltwater rearing conditions. Table 1 shows markers that remained statistically significant after an FDR correction. We considered linkage groups with more than one significant locus as being involved in adaptation to salinity. Linkage groups where heterozygotes were under-represented in fresh water and over-represented in salt water were: 3, 6, 7, 12, and 17 (Figure 2). The effects were particularly strong for linkage group 7 where the heterozygotes were 1.9 times as abundant in salt water (~0.65) as they were in fresh water (~0.34). Loci at linkage group 16 showed the opposite pattern where heterozygous individuals were common among freshwater and rare among saltwater offspring. Table S5 shows the results for all markers.

**Table 1.** Salinity associated loci.

| Marker | Linkage Group | Position (cM) | Proportion of heterozygotes in freshwater | Proportion of heterozygotes in saltwater | Chi-square P-value | FDR P-value |
| --- | --- | --- | --- | --- | --- | --- |
| 9418 | 7 | 16.96 | 0.34 | 0.65 | 4.76E-07 | 6.90E-05 |
| 141 | 7 | 17.49 | 0.34 | 0.64 | 1.31E-06 | 9.50E-05 |
| 14667 | 7 | 0.00 | 0.18 | 0.41 | 4.13E-06 | 0.0002 |
| 14398 | 17 | 24.83 | 0.44 | 0.67 | 0.0003 | 0.0095 |
| 18723 | 17 | 25.90 | 0.45 | 0.67 | 0.0006 | 0.0171 |
| 11877 | 21 | 0.00 | 0.54 | 0.33 | 0.0007 | 0.0175 |
| 137 | 17 | 25.89 | 0.46 | 0.66 | 0.0014 | 0.0293 |
| 13073 | 6 | 0.00 | 0.48 | 0.67 | 0.0018 | 0.0331 |
| 13872 | 18 | 9.84 | 0.33 | 0.51 | 0.0022 | 0.0354 |
| 11937 | 3 | 0.00 | 0.34 | 0.52 | 0.0037 | 0.0472 |
| 10789 | 3 | 0.00 | 0.35 | 0.52 | 0.0048 | 0.0472 |
| 11514 | 3 | 0.08 | 0.34 | 0.51 | 0.0049 | 0.0472 |
| 14634 | 3 | 3.34 | 0.34 | 0.51 | 0.0059 | 0.0472 |
| 11023 | 12 | 6.11 | 0.41 | 0.58 | 0.0059 | 0.0472 |
| 15386 | 12 | 6.62 | 0.38 | 0.57 | 0.0036 | 0.0472 |
| 5062 | 12 | 6.66 | 0.39 | 0.57 | 0.0044 | 0.0472 |
| 10999 | 16 | 0.00 | 0.62 | 0.45 | 0.0043 | 0.0472 |
| 11538 | 16 | 3.18 | 0.61 | 0.43 | 0.0057 | 0.0472 |

**Figure 2.**
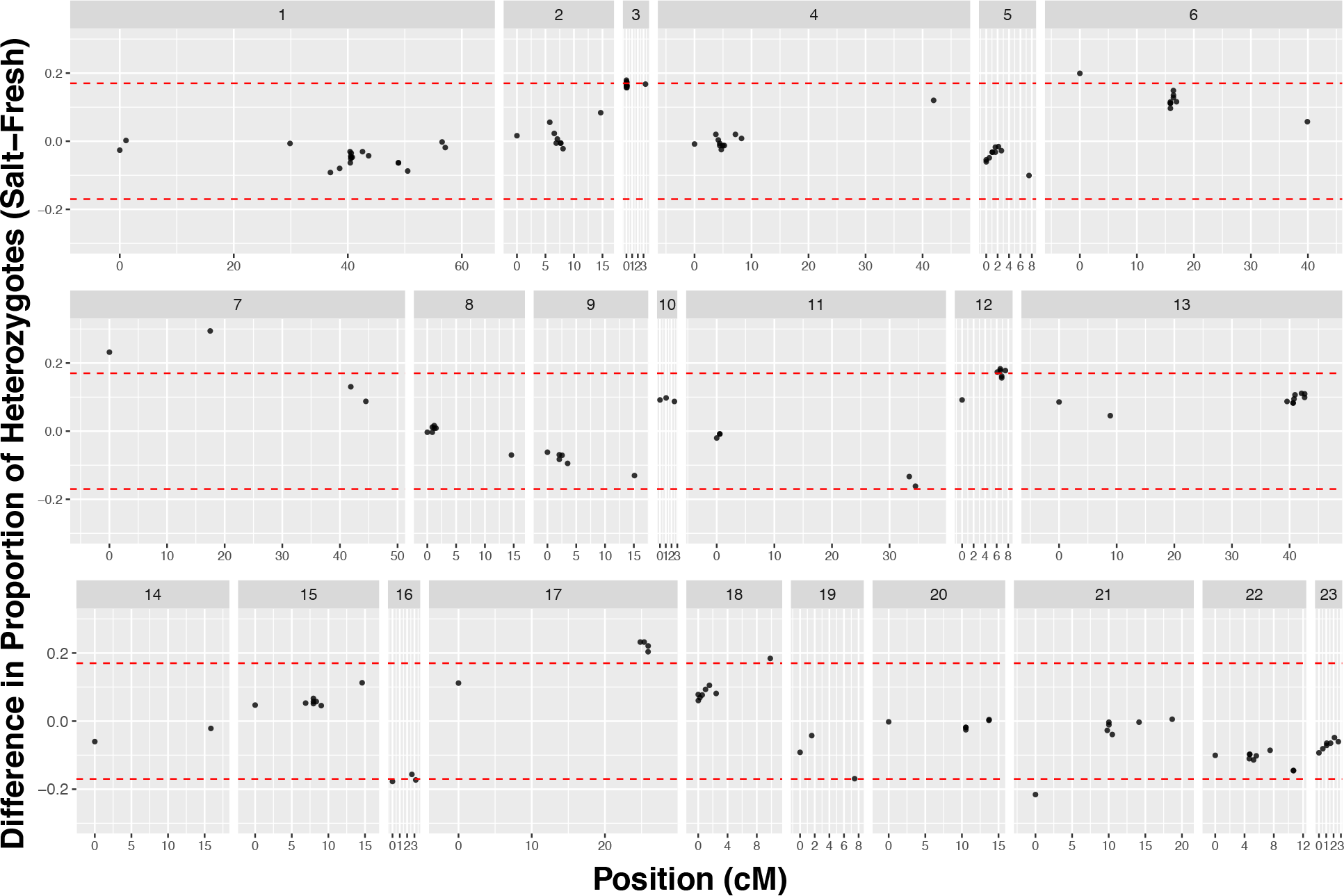
Salinity tolerance loci. Difference in proportion of heterozygous individuals in salt vs. freshwater plotted for loci across all 23 linkage groups. Linkage group numbers listed above, position of loci in centiMorgans (cM) on the hybrid map shown. (different linkage groups separated by white partitions). Red lines indicate FDR cutoffs. LG 3, 6, 7,12, 17 showed outliers. See Table 1 for loci names.

### Fertility and Hatching success as a Function of Male Genotype

Male fertility (proportion of unfertilized eggs) mapped to a single QTL located on linkage group 7 at 25 cM (LOD= 4.15, p = 0.034, Figure 3A). Hybrid viability, the proportion of fertilized eggs surviving to hatching, mapped to linkage group 1 at 9 cM (LOD = 3.47, p = 0.038; Figure 3B).

**Figure 3.**
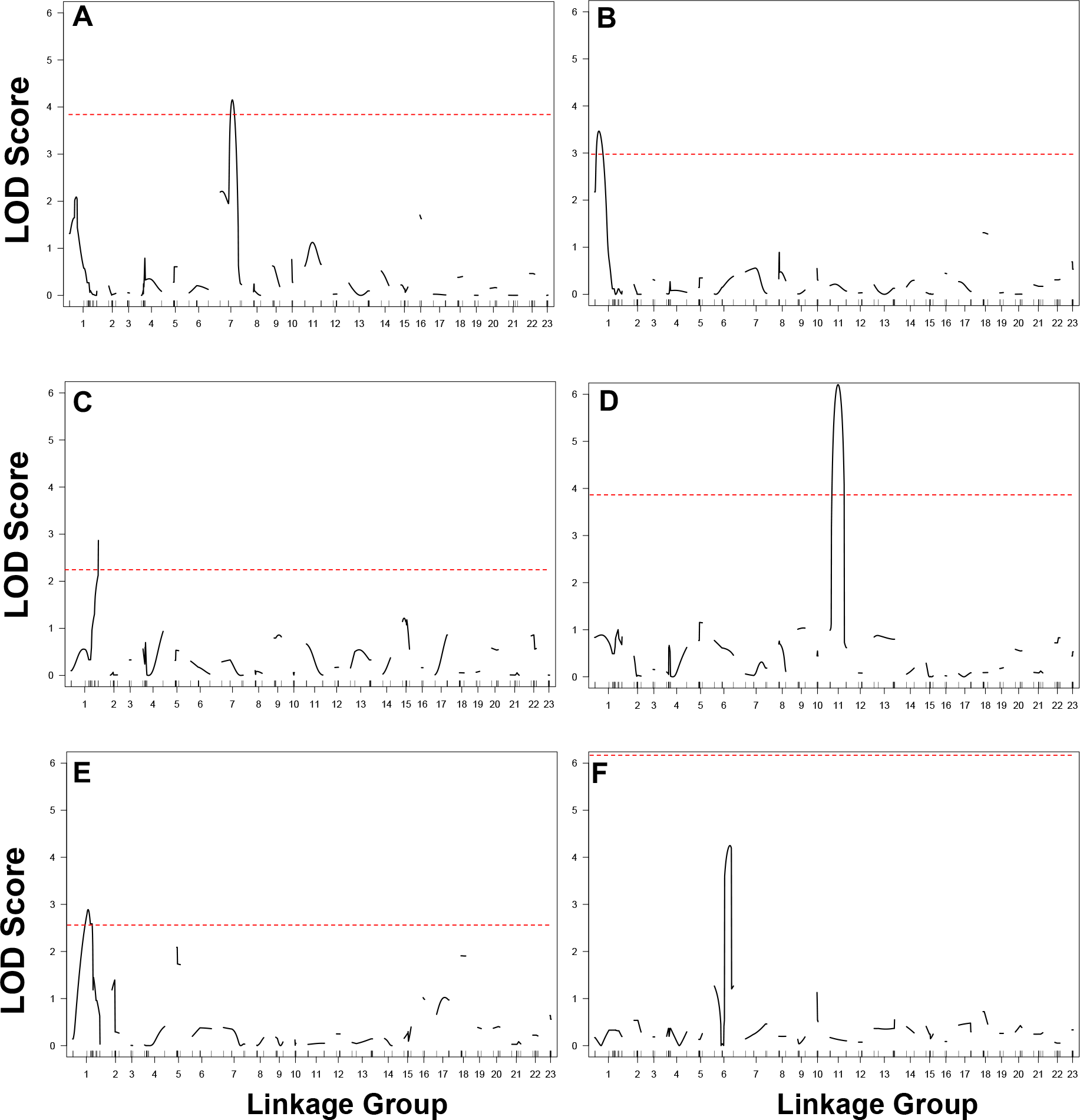
LOD scores from QTL mapping of reproductive isolating barriers. A) Male fertility, B) male offspring hatching success, C) male probability of mating with *L. parva,* D) number of eggs produced when male mated to *L. parva*, E) number of eggs produced when male mated to *L. goodei,* F) number of eggs produced when females are mated *L. parva .* Red dashed line indicates the p < 0.05 threshold determined by permutation, which varies due to differences in number of phenotyped individuals and whether or not a binary model is used for each trait.

### Behavioral isolation

For backcrossed males, the probability of mating occurring over 20 days was only 52% when paired with *L. parva* females and this trait mapped to linkage group 1, marker 13870 at 57 cM (LOD=2.87, p=0.028; Figure 3C). Males heterozygous for the *L. parva* allele at chromosome 1 were less likely to mate with *L. parva*, suggesting that this allele may not confer attractiveness and may represent an incompatibility. There were no QTL identified for the probability of a male mating with *L. goodei* females. The number of eggs laid when males were mated to *L. parva* females mapped to chromosome 11 at 16.5 cM (LOD = 6.2, p = 0.004) (Figure 3D). The number of eggs a male laid with a *L. goodei* female mapped to chromosome 1 at 32 cM (LOD = 2.89, p = 0.004; Figure 3E).

For backcrossed females, no significant QTL were identified. There was a weak association (p = 0.11) of number of eggs laid with *L. parva* males on chromosome 6 at 32.5 cM (LOD = 4.19; Figure 3F).

## DISCUSSION

In this study, we explored the role of extrinsic and intrinsic factors on speciation in the killifish *Lucania goodei* and *L. parva* by genetically mapping the sex determining locus, salinity tolerance, behavioral isolation, and hybrid incompatibilities. We found that salinity tolerance has a polygenic basis but adaptation to salinity is unlikely to have contributed strongly to the development of reproductive isolation in this system as salinity tolerance loci rarely overlap with isolating loci. Instead, a fusion between the chromosome with the sex determining locus and an autosome in *L. parva* appears to have significantly contributed to speciation as multiple different components of reproductive isolation mapped there (Figure 4). Below we discuss these results in more detail.

Salinity tolerance mapped to numerous locations in the *Lucania* genome revealing a strong polygenic basis to this trait. This is not surprising as decades of research have revealed that salinity tolerance in teleosts is a complex trait that involves multiple tissues (e.g, gills, kidneys) and physiological pathways (Evans *et al.*, 2005; Evans, 2008; Larsen *et al.*, 2011; Laverty & Skadhauge, 2012). We found that the loci underlying this trait were not grouped together in a single area but were instead spread out across the genome. Other studies of the genomic basis of salinity tolerance in teleosts have revealed similarly distributed genetic architectures. For example, a comparison of salinity tolerance QTL in three different salmonids revealed that between 3 (in *Oncorhynchus mykiss*) and 10 (*Salmo salar* and *Salvelinus alpinus*) linkage groups are involved (Norman *et al.*, 2012). Salinity tolerance in Atlantic cod (*Gadus morhua)* maps to 11 different linkage groups (Berg *et al.*, 2015). It is unclear if this kind of genetic architecture will facilitate or hinder the development of reproductive isolation in speciation with gene flow. For instance, it will be difficult to maintain linkage disequilibrium between loci that are spread out over many linkage groups when gene flow is high. However, spreading divergent selection across the genome increases the chance for processes such as divergence hitchiking (Via & West, 2008; Via, 2009), leading to increased genome divergence overall.

**Figure 4.**
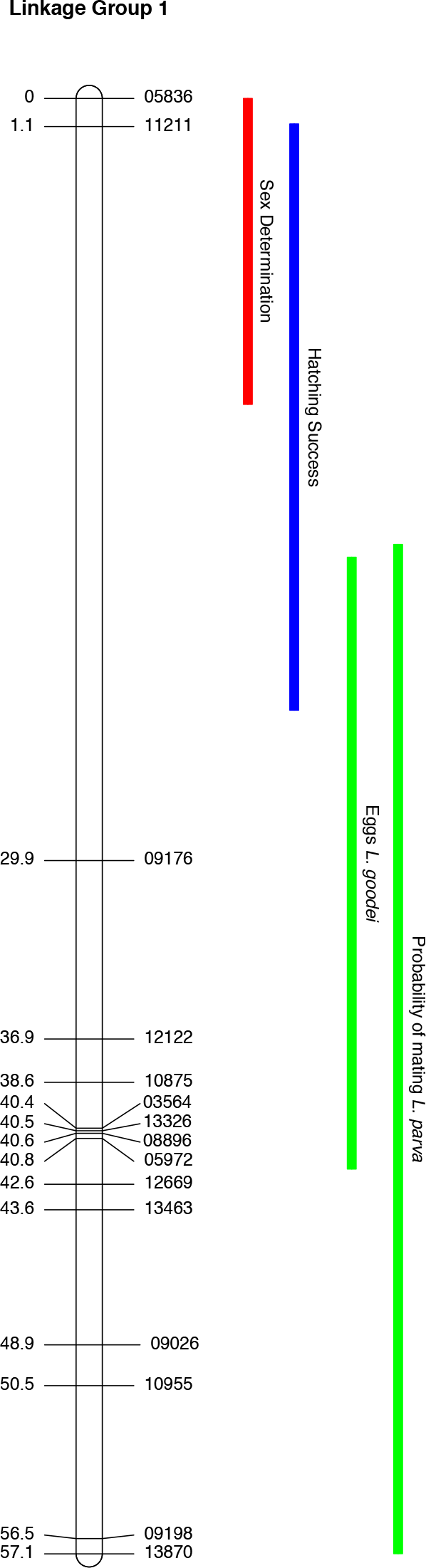
Sex determining and isolating loci mapping to Linkage group 1. Bayesian credible intervals for sex determination and isolating traits (solid rectangles) mapped relative to position (in cM) along linkage group 1 (the fused chromosome) from the hybrid linkage map. Blue indicates hybrid incompatibility; green indicates behavioral isolation; sex locus shown in red. The ancestral autosomal portion is ~40-57 cM on this hybrid map.

We found little evidence in this study that divergent selection for salinity tolerance in *Lucania* actually generated reproductive isolation. There are several different ways that divergent natural selection may generate reproductive isolation. The majority of these mechanisms, such as magic traits (Gavrilets 2004) and divergence hitchiking (Via & West, 2008, Via, 2009), predict that traits that are under divergent natural selection and those that contribute to reproductive isolation map to the same area of the genome. Although salinity tolerance mapped to 4 different linkage groups, only linkage group 7 also contained a locus involved in reproductive isolation, contributing to male fertility. When backcrossed individuals carry *L. goodei* alleles on linkage group 7, they were more likely to be infertile and survive poorly at high salinities. This area of the genome is interesting because genomic scans suggest that *L. goodei* and *L. parva* are differentiated in both sperm-related and ion transport genes (Kozak *et al.,* 2014). However, we did not detect enough overall co-localization to implicate a general role for natural selection to salinity leading to divergence hitchhiking or multiple reproductive barriers. There are other mechanisms by which natural selection may lead to reproductive isolation without the co-localization of loci. For example, sensory bias may have led to sexual signals that are strongly adapted to different salinity environments. However, this mechanism has already been ruled out in this system (Berdan & Fuller, 2012). Thus, divergent natural selection is unlikely to have directly contributed to the evolution of reproductive isolation in *Lucania* killifish.

The chromosomal fusion seems to have played a significant role in the speciation between *L. goodei* and *L. parva* as several components of reproductive isolation map there (Figure 4; Table 3). The male sex determining loci mapped to the fused chromosome in both hybrid (*L. goodei* × *L. parva*) and pure *L. parva* crosses. This suggests that this Robertsonian fusion in *L. parva* occurred between the *Lucania* chromosome with the sex determining loci and an autosome. Chromosomal fusions, often differentiate populations or species and have been shown both theoretically and empirically to facilitate adaptation (Franchini *et al.*, 2010; Guerrero & Kirkpatrick, 2014; Dobigny *et al.*, 2017; Wellband *et al.*, 2019). In fishes, sex chromosomes are often involved in fusions possibly because fusions resolve sexually antagonistic selection (Kitano & Peichel, 2012) or because male-mutation bias leads to Y-fusions (Pennell *et al*., 2015). However, unlike many other known fusions in fish, our fusion does not appear to represent a neo-Y system with unfused X chromosomes, because both males and females possess fused chromosomes (Berdan *et al.*, 2014). Our results add to the growing evidence that chromosomal fusions may facilitate evolutionary processes.

**Table 3.**
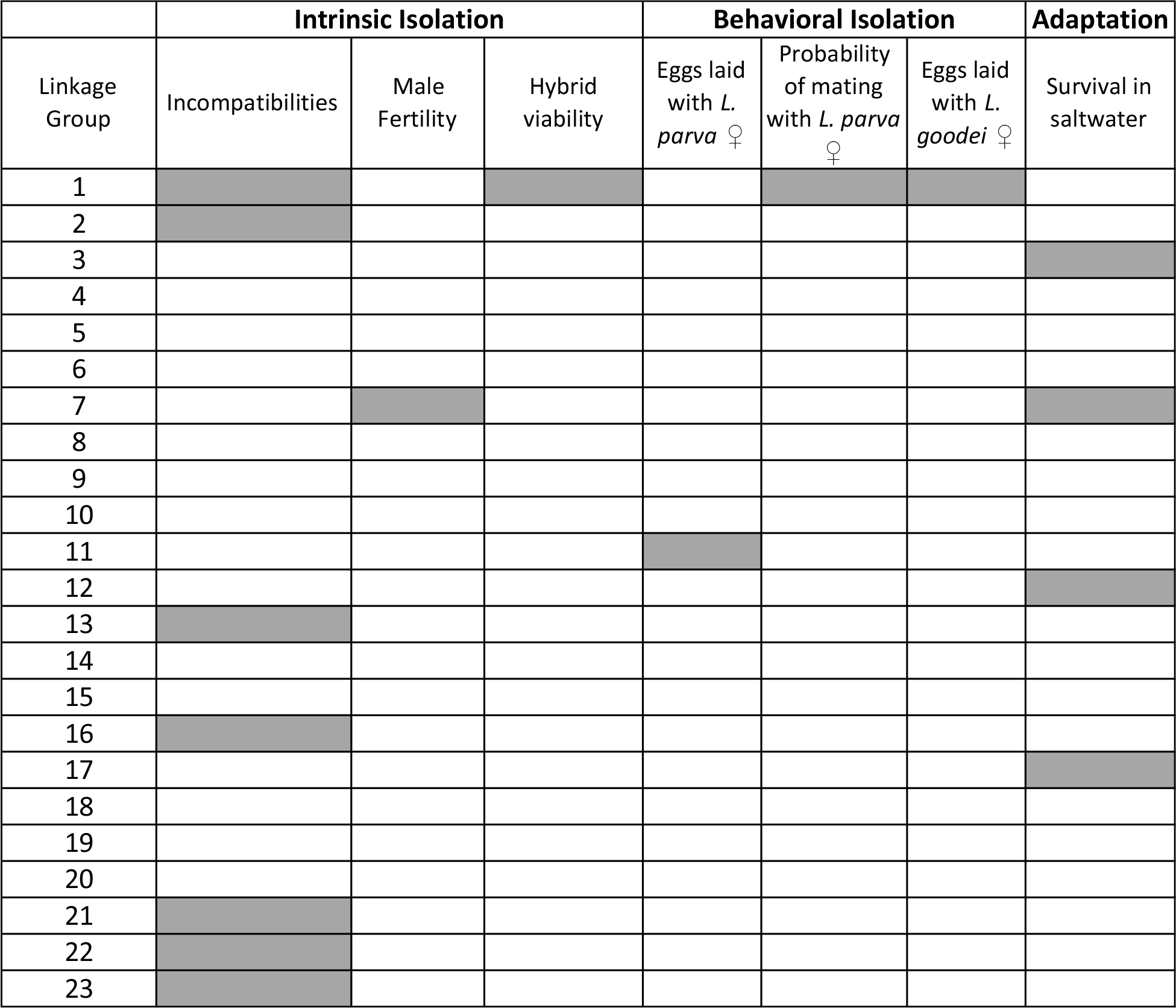
Summary of locations of isolating barriers.

We found that the fused chromosome contained QTLs for both behavioral isolation (number of eggs laid with *L. goodei*, probability of mating with *L. parva* females) and hybrid incompatibilities (number of offspring that survived to hatching and loci contributing to gametic disequilibrium). Only one other linkage group (LG 7) contained more than a single trait, with salinity and fertility mapping to LG 7. In order for speciation with gene flow to proceed, different forms to reproductive isolation must be coupled with one another (Smadja & Butlin, 2011; Butlin & Smadja, 2018). Physical linkage/reduced recombination is one of the strongest ways to generate linkage disequilibrium and chromosomal rearrangements often play a role in generating this reduced recombination (Hoffmann & Rieseberg, 2008; Faria & Navarro, 2010; Wellenreuther & Bernatchez, 2018; Wellenreuther *et al.*, 2019). Chromosomal fusions can generate linkage disequilibrium in two ways: first by bringing previously unlinked loci together and second by reducing recombination, especially around the centromere (Dumas & Britton-Davidian, 2002; Franchini *et al.*, 2010). However, only the QTL for probability of mating with *L. parva* mapped to the formerly autosomal portion of the chromosome (~40-57 cM) and this locus appeared to function as an incompatibility, with the *L. parva* alleles in an *L. goodei* background contributing to low mating success. Future work will be needed to determine if physical linkage of this locus with the other isolating loci was a benefit provided by the fusion, similar to the situation in Japan Sea sticklebacks where the Y-chromosome fused to an autosome containing a behavioral isolation locus (Kitano *et al.*, 2009).

The genetic architecture of reproductive isolation in *Lucania* is conducive to the process of reinforcement. Reinforcement occurs when hybrids suffer reduced fitness which generates selection for increased behavioral isolation in areas of sympatry to avoid mating with heterospecifics (Servedio & Noor, 2003). Previous behavioral work has found that reinforcement has contributed significantly to the evolution of species-specific preferences in sympatry in both sexes of *L. goodei* and *L. parva* (Gregorio *et al.*, 2012; Kozak *et al.*, 2015). Theoretical studies show stronger reinforcement when incompatibility loci and loci for behavioral isolation are linked to sex than when they are located on autosomes (Servedio & Saetre, 2003; Lemmon & Kirkpatrick, 2006; Hall & Kirkpatrick, 2006). The co-localization of both behavioral and incompatibility loci to the sex chromosome we find is consistent with this theory and the known role of reinforcement in driving speciation in *Lucania.* Indeed, the degree of sex-linkage of isolating loci may have predisposed *Lucania* mate preferences toward rapid evolution in sympatry.

In summary, we find that the fused sex chromosome in *L. parva* contributes disproportionately to reproductive isolation between *L. parva* and *L. goodei*. Salinity tolerance in *L. parva* is polygenic, distributed across the genome, and rarely co-localizes with reproductive isolating traits. Speciation in this system appears to be driven by reinforcement and indirect selection against hybrids rather than direct natural selection for salinity tolerance.

## ACKNOWLEDGEMENTS

This work was funded by a National Science Foundation (NSF) Early Faculty Development (CAREER) grant (DEB-0953716) to R.C.F, a National Science Foundation Dissertation Improvement Grant to R.C.F and E.L.B. (DEB-1110658), and a University of Illinois Research Board Award 11153. This work was approved by the University of Illinois Institutional Animal Care and Use Committee (protocols 08183 and 09306).

