## Supplemental Figures for "Genomic landscape of reproductive isolation in *Lucania* killifish: The role of sex chromosomes and salinity"

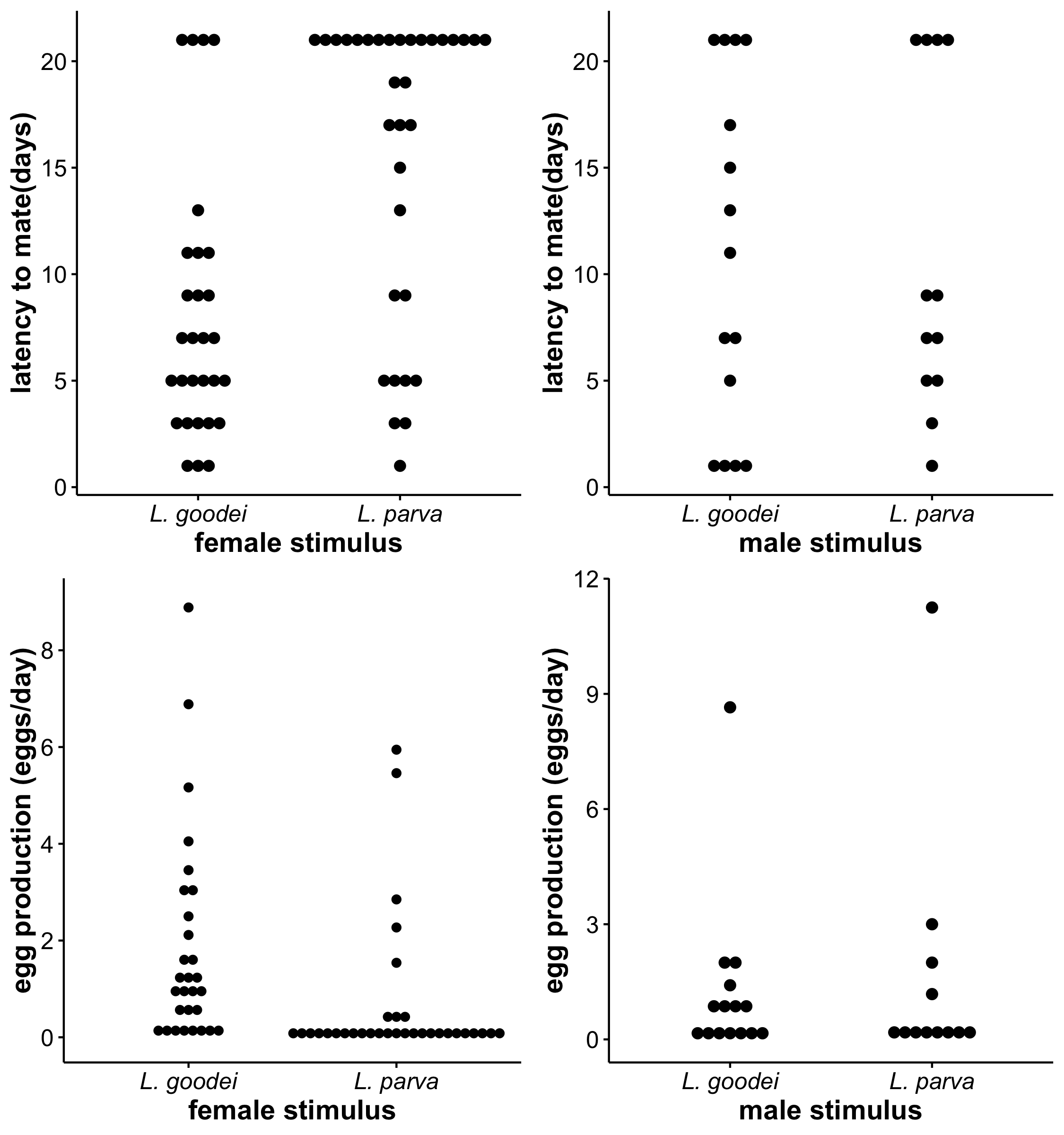

Supplemental Figure 1 - Raw data from mate choice trials. Data from (clockwise starting with top right): Female mating preference - latency to mate, Female mating preference - egg production, Male mating preference - egg production, and Male mating preference - latency to mate. Each dot represents a single backcrossed male or female.
